# Generative language modeling for antibody design

**DOI:** 10.1101/2021.12.13.472419

**Authors:** Richard W. Shuai, Jeffrey A. Ruffolo, Jeffrey J. Gray

**Affiliations:** Department of Electrical Engineering and Computer Sciences, University of California, Berkeley, CA; Program in Molecular Biophysics, The Johns Hopkins University, Baltimore, MD; Department of Chemical and Biomolecular Engineering, The Johns Hopkins University, Baltimore, MD

**Keywords:** antibodies, deep learning, language modeling

## Abstract

Discovery and optimization of monoclonal antibodies for therapeutic applications relies on large sequence libraries, but is hindered by developability issues such as low solubility, low thermal stability, high aggregation, and high immunogenicity. Generative language models, trained on millions of protein sequences, are a powerful tool for on-demand generation of realistic, diverse sequences. We present Immunoglobulin Language Model (IgLM), a deep generative language model for creating synthetic libraries by re-designing variable-length spans of antibody sequences. IgLM formulates antibody design as an autoregressive sequence generation task based on text-infilling in natural language. We trained IgLM on 558M antibody heavy- and light-chain variable sequences, conditioning on each sequence’s chain type and species-of-origin. We demonstrate that IgLM can generate full-length heavy and light chain sequences from a variety of species, as well as infilled CDR loop libraries with improved developability profiles. IgLM is a powerful tool for antibody design and should be useful in a variety of applications.

## Introduction

Antibodies have become popular for therapeutics because of their diversity and ability to bind antigens with high specificity (1). Traditionally, monoclonal antibodies (mAbs) have been obtained using hybridoma technology, which requires the immunization of animals (2). In 1985, the development of phage display technology allowed for in vitro selection of specific, high-affinity mAbs from large antibody libraries (3–5). Despite such advances, therapeutic mAbs derived from display technologies face issues with developability, such as poor expression, low solubility, low thermal stability, and high aggregation (6, 7). Display technologies rely on a high-quality and diverse antibody library as a starting point to isolate high-affinity antibodies that are more developable (8). Synthetic antibody libraries are prepared by introducing synthetic DNA into regions of the antibody sequences that define the complementarity-determining regions (CDRs), allowing for human-made antigen-binding sites. However, the space of possible synthetic antibody sequences is very large (diversifying 10 positions of a CDR yields 20^10^ ≈ 10^13^ possible variants). To discover antibodies with high affinity, massive synthetic libraries on the order of 10^10^–10^11^ variants must be constructed, often containing substantial fractions of non-functional antibodies (2, 8).

Recent work has leveraged natural language processing methods for unsupervised pre-training on massive databases of raw protein sequences for which structural data is unavailable (9–11). These works have explored a variety of pre-training tasks and downstream model applications. For example, the ESM family of models (trained for masked language modeling) have been applied to representation learning (9), variant effect prediction (12), and protein structure prediction (13). Autoregressive language modeling, an alternative paradigm for pre-training, has also been applied to protein sequence modeling. Such models have been shown to generate diverse protein sequences, which often adopt natural folds despite diverging signifcantly in residue makeup (14, 15). In some cases, these generated sequences even retain enzymatic activity comparable to natural proteins (16). Autoregressive language models have also been shown to be powerful zero-shot predictors of protein fitness, with performance in some cases continuing to improve with model scale (15, 17).

Another set of language models have been developed specifically for antibody-related tasks. The majority of prior work in this area has focused on masked language modeling of sequences in the Observed Antibody Space (OAS) database (18). Prihoda et al. developed Sapiens, a pair of distinct models (each with 569K parameters) for heavy and light chain masked language modeling (19). The Sapiens models were trained on 20M and 19M heavy and light chains respectively, and shown to be effective tools for antibody humanization. Ruffolo et al. developed AntiBERTy, a single masked language model (26M parameters) trained on a corpus of 558M sequences, including both heavy and light chains (20). AntiBERTy has been applied to representation learning for protein structure prediction (21). Leem et al. developed AntiBERTa, a single masked language model (86M parameters) trained on a corpus of 67M antibody sequences (both heavy and light). Representations for AntiBERTa were used for paratope prediction. Olsen et al. developed AbLang, a pair of masked language models trained on 14M heavy chains and 187K light chains, for sequence restoration (22). For sequence generation, autoregressive generative models have been trained on antibody sequences and used for library design (23, 24). Akbar et al. (23) trained an LSTM for autoregressive generation of CDR H3 loops and conducted an in silico investigation of their potential for binding antigens. Shin et al. (24) experimentally validated a set of nanobody sequences with generated CDR3 loops and showed promising improvements to viability and binding discovery when compared to traditional approaches, despite the library being over 1000-fold smaller. However, because this generative model was unidirectional, it could not be used to directly re-design the CDR3 loop *within the sequence*, and instead had to be oversampled to produce sequences matching the residues following the loop.

Here, we present Immunoglobulin Language Model (IgLM), a generative language model that leverages bidirectional context for designing antibody sequence spans of varying lengths while training on a large-scale natural antibody dataset. We show that IgLM can generate full-length antibody sequences conditioned on chain type and species-of-origin. Furthermore, IgLM can diversify loops on an antibody to generate high-quality libraries that display favorable biophysical properties while resembling human antibodies. IgLM should be a powerful tool for antibody discovery and optimization.

## Results

### Immunoglobulin language model

Our method for antibody sequence generation, IgLM, is trained on 558 million natural antibody sequences for both targeted infilling of residue spans, as well as full-length sequence generation. IgLM generates sequences conditioned on the species-of-interest and chain type (heavy or light), enabling controllable generation of antibody sequences.

#### Infilling language model

Design of antibody libraries typically focuses on diversification of the CDR loop sequences in order to facilitate binding to a diverse set of antigens. Existing approaches to protein sequence generation (including antibodies) typically adopt left-to-right decoding strategies. While these models have proven effective for generation of diverse and functional sequences, they are ill-equipped to re-design specific segments of interest within proteins. To address this limitation, we developed IgLM, an infilling language model for immunoglobulin sequences. IgLM utilizes a standard left-to-right decoder-only transformer architecture (GPT-2), but it is trained for infilling through rearrangement of sequences. Specifically, we adopt the infilling language model formulation from natural language processing (25), wherein arbitrary-length sequence segments (spans) are masked during training and appended to the end of the sequence. By training on these rearranged sequences, models learn to predict the masked spans conditioned on the surrounding sequence context.

To train IgLM, we collected antibody sequences from the Observed Antibody Space (OAS) (18). The OAS database contains natural antibody sequences from six species: human, mouse, rat, rabbit, rhesus, and camel. To investigate the impacts of model capacity, we trained two versions of the model: IgLM and IgLM-S, with 13M and 1.4M trainable parameters, respectively. Both IgLM models were trained on a set of 558M non-redundant sequences, clustered at 95% sequence identity. During training, we randomly masked spans of ten to twenty residues within the antibody sequence to enable diversification of arbitrary spans during inference. Additionally, we conditioned sequences on the chain type (heavy or light) and species-of-origin. Providing this context enables controllable generation of species-specific antibody sequences. An example of training data construction is illustrated in Figure 1A. Unless otherwise specified, we use the larger IgLM model for all experiments.

**Fig. 1.**
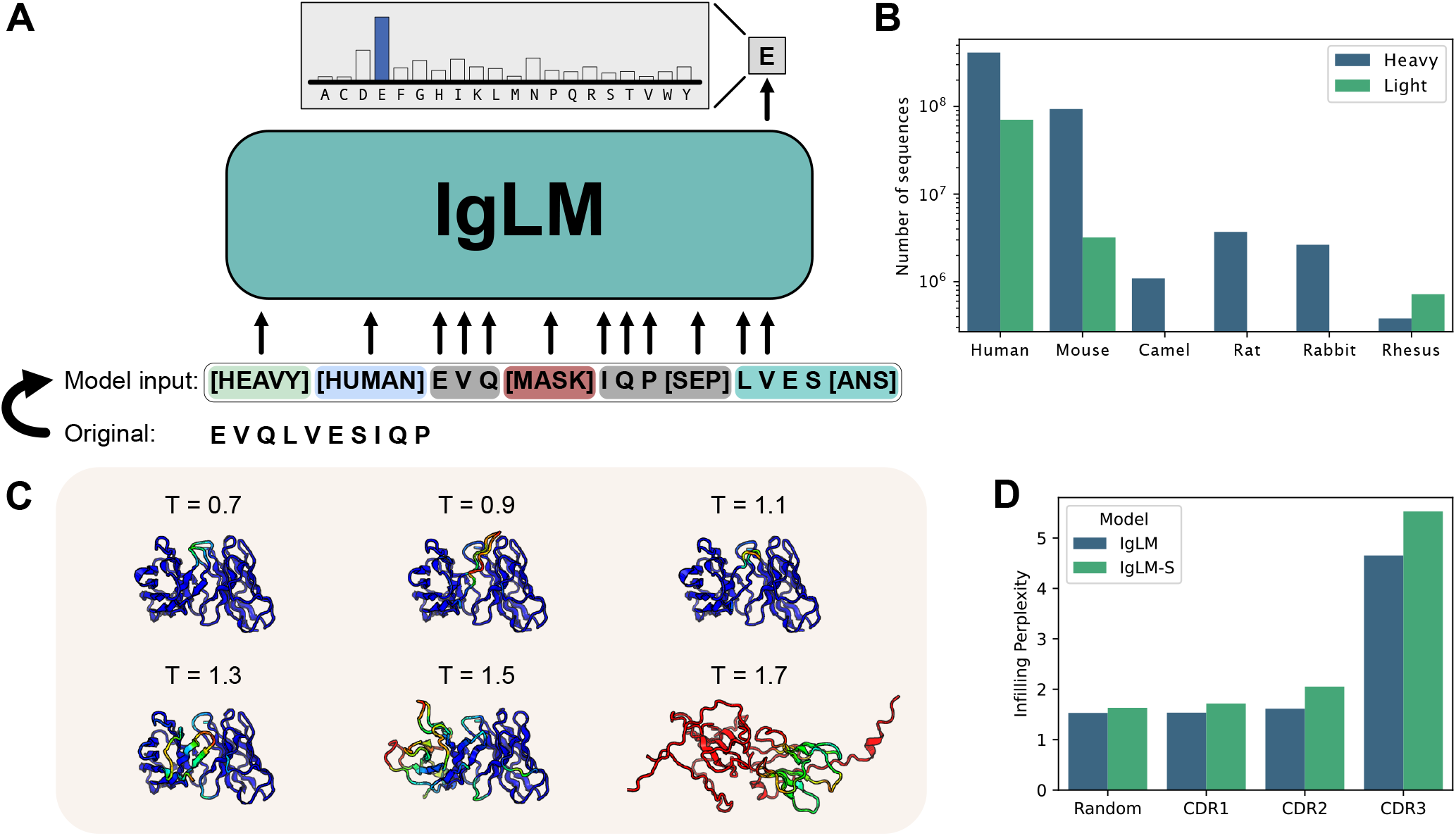
Overview of IgLM model for antibody sequence generation. (A) IgLM is trained by autoregressive language modeling of reordered antibody sequence segments, conditioned on chain and species identifier tags. (B) Distribution of sequences in clustered OAS dataset for various species and chain types. (C) Infilling perplexity for IgLM and IgLM-S on heldout test dataset for CDR loops and random spans of 10-20 residues within sequences. (D) Effect of increased sampling temperature for full-length generation. Structures at each temperature are predicted by AlphaFold-Multimer and colored by prediction confidence (pLDDT), with blue being the most confident and red being the least.

#### IgLM generates foldable antibody sequences

As an initial validation of the antibody sequence generation capabilities of IgLM, we conducted a small scale investigation of full-length generation. Specifically, we investigated the impacts of sampling temperature for tuning the diversity of generated sequences. Sampling temperature values above one effectively flatten the amino acid distribution at each step of generation, resulting in more diverse sequences, while temperature below one sharpens the distribution at each position, resembling a greedy decoding strategy. We generated a set of full-length sequences at temperatures ranging from 0.7 to 1.7, providing the model with human heavy and human light conditioning tags. Because IgLM was trained for sequence infilling, generated sequences contain discontinuous segments of sequence segments, which we simply reordered to produce full-length antibodies. Generated heavy and light chain sequences were paired according to sampling temperature and their structures were predicted using AlphaFold-Multimer (26). In general, IgLM generates sequences with correspondingly confident predicted structures at lower temperatures (up to 1.3), before beginning to degrade in quality at higher temperatures (Figure 1C).

#### Language modeling evaluation

We evaluated IgLM as a language model by computing the per-token perplexity for infilled spans within an antibody, which we term the *infilling perplexity*. Because the infilled segment is located at the end of the sequences, computing the infilling perplexity is equivalent to taking the per-token perplexity after the [`SEP`] token. We compared the infilling perplexity of IgLM and IgLM-S on a heldout test dataset of 30M sequences (Figure 1D). Results are tabulated by CDR loop, as well as for spans selected randomly within the antibody sequence. As expected, we observe greater perplexity for the CDR loops than the randomly chosen spans, which include the highly conserved framework regions. The CDR3 loop, which is the longest and most diverse, has the highest infilling perplexity. When we compare IgLM and IgLM-S, we observe that IgLM has a lower infilling perplexity for all CDR loops, indicating that the larger IgLM model (with ten times more parameters) is better at modeling the diversity of antibody sequences.

The diversity of antibody sequences varies by species and chain type. For example, heavy chains introduce additional diversity into their CDR3 loops via D-genes, while some species (e.g., camels) tend to have longer loops. To investigate how these differences impact the performance of IgLM in different settings, we also tabulated the heldout set infilling perplexity by species and chain type. In general, both IgLM models achieve low infilling perplexity for random spans across all species (Figure S1). For CDR1 and CDR2 loop infilling, perplexity values are typically lower for human and mouse antibodies, which are disproportionately represented in the OAS database. In general, both models still perform better on these loops than the more challenging CDR3 loops, regardless of species. One exception is for rhesus CDR2 loops, on which IgLM-S performs considerably worse than the larger IgLM model. This appears to be due to poor fitting of rhesus CDR L2 loops, as reflected in the similarity high infilling average perplexity observed when tabulated by chain type (Figure S2). The highest infilling perplexity is observed for camel CDR3 loops, which tend to be longer than other species. Across all species and chain types, the larger IgLM model achieves lower infilling perplexity than IgLM-S, suggesting that further increasing model capacity would yield additional improvements.

### Controllable generation of antibody sequences

Having demonstrated that IgLM can generate well-formed full-length sequences, we next considered the controllability of IgLM for generating antibody sequences with specific traits. Controllable generation utilizes conditioning tags to provide the model with additional context about the expected sequence.

#### Generating species- and chain-controlled sequences

To evaluate the controllability of IgLM, we generated a set of 220K full-length sequences utilizing all viable combinations of conditioning tags, as well as a range of sampling temperatures (Figure 2A). For every species (except camel), we combined sampled with both heavy and light conditioning tags. For camel sequence generation, we only sampled heavy chains, as they do not produce light chains. To produce a diverse set of sequences for analysis, we sampled using a range of temperatures (*T* ∈ {0.6, 0.8,1.0,1.2}). Sampling under these conditions resulted in a diverse set of antibody sequences. However, we observed that the sequences frequently featured N-terminal truncations, a common occurrence in the OAS database used for training (22). For heavy chains, these N-terminal deletions appeared as a left-shoulder in the sequence length distribution (Figure 2B, left) with lengths ranging from 100 to 110 residues. For light chains, we observed a population of truncated chains with lengths between 98 and 102 residues (Figure 2B, right). To address truncation in generated sequences, we utilized a prompting strategy, wherein we initialize each sequence with a three-residue motif corresponding to the species and chain type tags. Specific intialization sequences are documented in Table S2. For both heavy and light chains, prompting with initial reisdues significantly reduced the population of truncated sequences (Figure 2B). For the following analysis, we consider only sequences generated with prompting.

**Fig. 2.**
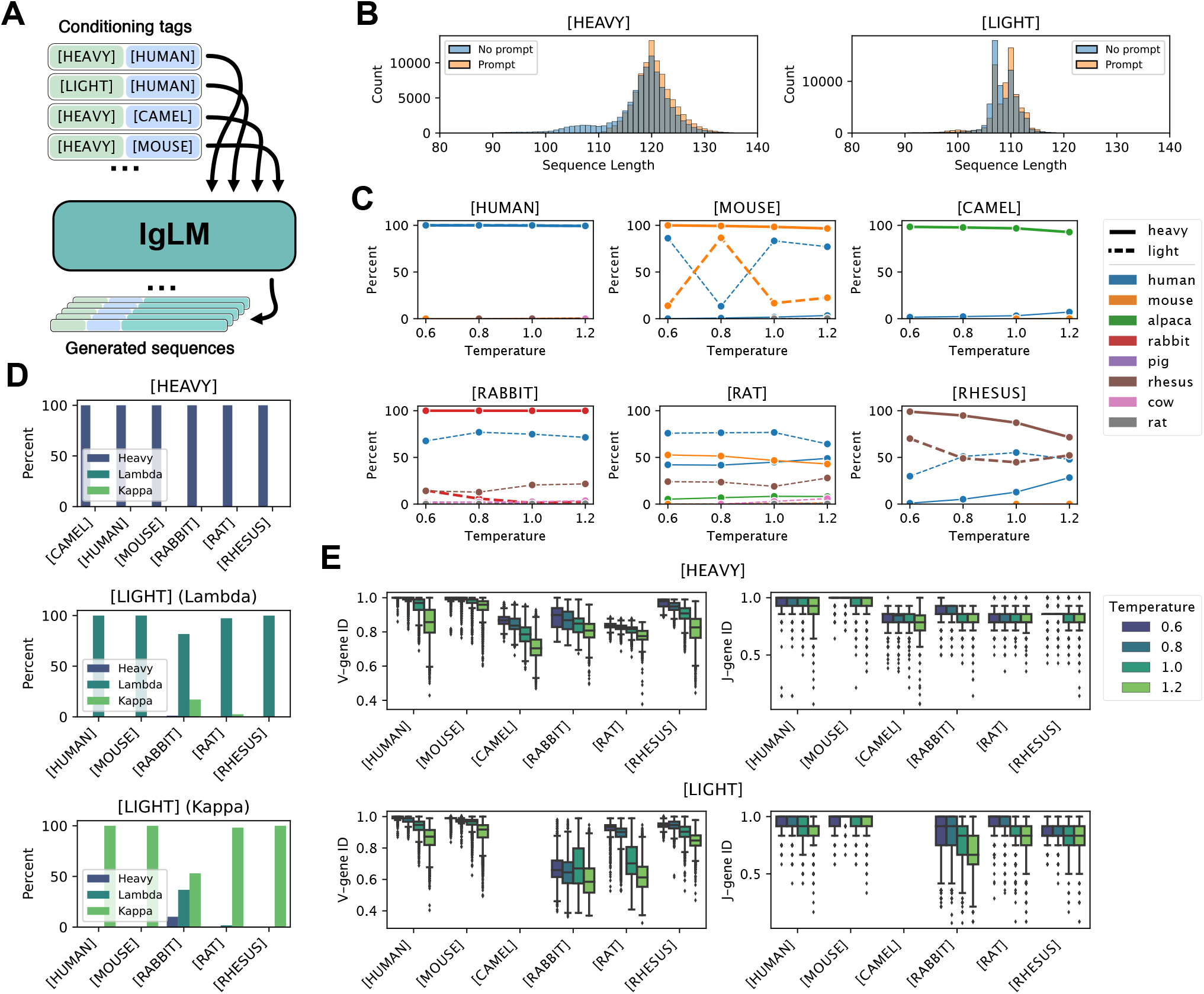
Controllable antibody sequence generation. (A) Diagram of procedure for generating full-length antibody sequences given a desired species and chain type with IgLM. (B) Length of generated heeavy and light with and without initial three residues provided (prompting). (C) Adherence of generated sequences to species conditioning tags. Each plot shows the species classifications of antibody sequences generated with a particular species conditioning tag (indicated above plots). Solid and dashed lines correspond to sequences generated with heavy- and light-chain conditioning, respectively. (D) Adherence of generated sequences to chain conditioning tags. Top plot shows the percentage of heavy-chain-conditioned sequences classified as heavy chains, for each species conditioning tag. Lower plots show the percentage of light-chain-conditioned sequences, further divided by whether initial residues were characteristic of lambda or kappa chains, classified as lambda or kappa chains. (E) Effect of sampling temperature on germline identity for generated heavy and light chain sequences. As sampling temperature increases, generated sequences diverge from the closest germline V- and J-gene sequences.

#### Adherence to conditioning tags

To evaluate the effectiveness of contrallable generation, we considered the agreement between the provided conditioning tags and the sequences produced by IgLM. For each generated sequence, we classified the species (according to V-gene identity) and chain type using ANARCI (27). We note that the species classes provided by ANARCI diverge in some cases from those provided by the OAS database, but there was a suitable corresponding class for each conditioning token (e.g., alpaca for [`CAMEL`]). In Figure 2C, we show the makeup of sequences for each species conditioning tag, according to sampling temperature. In each plot, the percentage of heavy and light chain sequences classified as each species are indicated by solid and dashed lines, respectively. For most species (human, mouse, camel, rabbit, rhesus), IgLM is able to successfully generate heavy chain sequences at every temperature. The exception to this trend is rat sequences, for which we were unable to produce any sequences that ANARCI classified as belonging to the intended species.

The ability to generate sequences is not directly explained by prevalence in the training dataset, as the model is trained on an order of magnitude more rat heavy chain sequences than rhesus (Table S1). IgLM is generally less effective at generating light chain sequences for most species. With the exception of human light chains, all species have a large proportion of sequences classified as belonging to an unintended species (typically human). For mouse and rhesus light chains, IgLM generates the correct species in 34.89% and 88.14% of cases, respectively (Table S3). For rabbit and rat light chains, IgLM was not exposed to any examples during training. Interestingly, despite having seen no such sequences during training, IgLM is capable of generating sequences classified by ANARCI as rabbit light chains for 6.89% of samples (1,120 sequences). The majority of these sequences are cases where the model has instead generated a rabbit heavy chain. However, for 35 of these 1,120 cases, IgLM has produced rabbit light chain sequences. We further investigated the plausibility of these sequences by aligning to the nearest germline sequences assigned by ANARCI with Clustal-Omega (28). The sequences appear to align well to rabbit germlines, though with conderable mutations to the framework regions (Figure S3). To investigate the structural viability of the generated rabbit light chain sequences, we predicted structures with IgFold (21). All structures were predicted confidently in the framework residues, with the CDR loops being the most uncertain (Figure S4). Although rare (35 sequences out of 20,000 attempts), these results suggest that IgLM is capable of generating rabbit light chain sequences despite having never observed such sequences during training. This may be achieved by producing a consensus light chain, with some rabbit-likeness conferred from the heavy chain examples.

We next evaluated the adherence of IgLM-generated sequences to chain type conditioning tags. In Figure 2D, we show the percentage of sequences classifed by ANARCI as heavy or light for each conditioning tag. Light chains are further divided into lambda and kappa classes. When conditioned towards heavy chain generation, IgLM effectively produces heavy chains for all species. For light chains, we observe a similar trend, with IgLM producing predominantly light chain sequences for all species. Only for rabbit sequences do we observe a population of heavy chains when conditioning for light chains. As noted above, these are cases where IgLM has instead produced a rabbit heavy chain. When generating light chain sequences, we provide initial residues characteristic of both lambda and kappa chains in equal proportion (Figure S2). For most species (except rabbit), the generated sequences are aligned with light chain type indicated by the initial residues. However, as noted above, many of the light sequences for poorly represented species are human-like, rather than resembling the desired species. Interestingly, these results suggest that the chain type conditioning tag is a more effective prior for IgLM than species.

#### Sampling temperature controls mutational load

Increasing sampling temperature has the effect of flattening the probability distribution at each position during sampling, resulting in a greater diversity of sequences. We evaluated the effect of sampling temperature on the diversity of generated sequences by measuring the fractional identity to the closest germline sequences using ANARCI (27). In Figure 2E, we show the germline identity for V- and J-genes for each species and chain type. At the lowest sampling temperature (*T* = 0.6), IgLM frequently recapitulates germline sequences in their entirety for some species (human, mouse, rhesus). As temperature increases, sequences for every species begin to diverge from germline, effectively acquiring mutations. Interestingly, J-gene sequences typically acquire fewer mutations than V-genes for both heavy and light chains. This is likely a reflection of the concentration of CDR loops within the V-gene (CDR1 and CDR2). Only a portion of the CDR3 loop is contributed by the J-gene, with the remaining sequence being conserved framework residues.

### Therapeutic antibody diversification

Diversification of antibody CDR loops is a common strategy for antibody discovery or optimization campaigns. Through infilling, IgLM is capable of replacing spans of amino acids within antibody sequences, conditioned on the surrounding context. To demonstrate this functionality, we generated infilled libraries for a set of therapeutic antibodies and evaluated several therapeutically relevant properties.

#### Infilled libraries for therapeutic antibodies

To evaluate the utility of infilling with IgLM for diversifying antibody sequences, we created infilled libraries for 49 therapeutic antibodies from Thera-SAbDab (29). For each antibody, we removed the CDR H3 loop (according to Chothia definitions (30)) and generated a library of infilled sequences using IgLM (Figure 3A). To produce diverse sequences, we used a combination of sampling temperatures (*T* ∈ {0.8,1.0,1.2}) and nucleus sampling probabilities (*P* ∈ {0.5, 0.75,1.0}). Nucleus sampling effectively clips the probability distribution at each position during sampling, such that only the most probable amino acids (summing to *P*) are considered. For each of the 49 therapeutic antibodies, we generated one thousand infilled sequences for each combination of *T* and *P*, totaling nine thousand variants per parent antibody. In Figure 3D, we show predicted structures (using IgFold (21)) for a subset of ten infilled loops derived from the trastuzumab antibody. The infilled loops vary in length and adopt distinct structural conformations. Across the infilled libraries, we see a variety of infilled CDR H3 loop lengths, dependent on the parent antibody’s surrounding sequence context (Figure 3B). The median length of infilled loops across antibodies ranges from 11 to 16 residues. Interestingly, we observe little impact on the length of infilled loops when varying the sampling temperature and nucleus probabilities (Figure 3C).

**Fig. 3.**
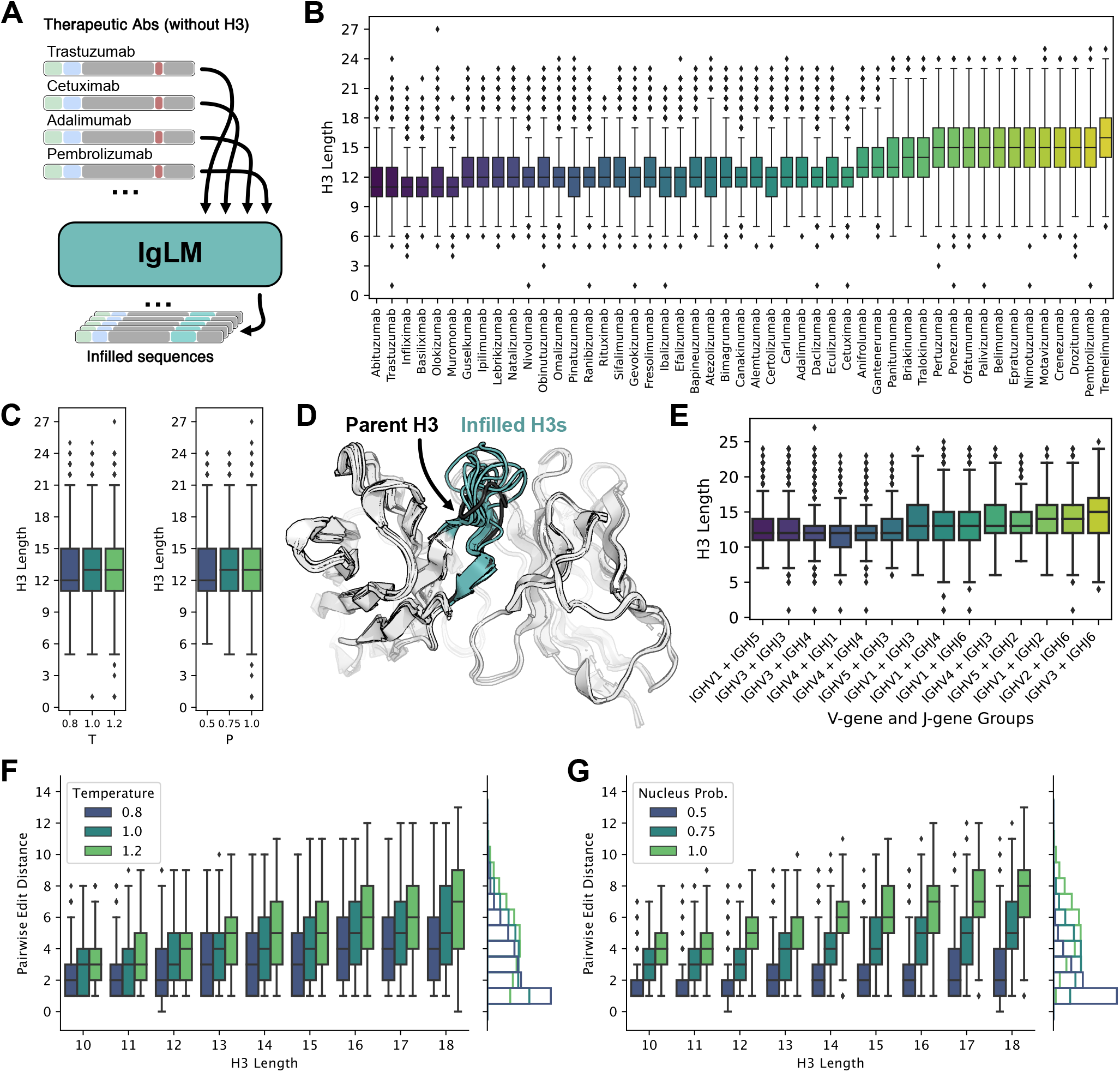
Generation of infilled therapeutic antibody libraries. (A) Diagram of procedure for generating diverse antibody libraries by infilling the CDR H3 loops of therapeutic antibodies. (B) Distribution of infilled CDR H3 loop lengths for 49 therapeutic antibodies. (C) Relationship between sampling temperature (*T*) and nucleus probability (*P*) and length of infilled CDR H3 loops. (D) Infilled CDR H3 loops for trastuzumab therapeutic antibody adopt diverse lengths and conformations. Structures for infilled variants are predicted with IgFold. (E) Distribution of infilled CDR H3 loop lengths for therapeutic antibodies grouped by nearest germline gene groups. (F-G) Effect of sampling temperature (*T*) and nucleus probability (*P*) on diversity of infilled CDR H3 loops for lengths between 10 and 18 residues. Pairwise edit distance measures the minimum edits between each infilled loop to another in the same set of generated sequences (i.e., within the set of sequences produced with the same *T* and *P* parameters). For both parameters, less restrictive sampling produces greater infilled loop diversity.

The distributions of infilled loop lengths vary considerably over the 49 therapeutic antibodies. Because IgLM is trained on natural antibody sequences, we hypothesized that the model may be performing a sort of germline matching, wherein sequences with similar V- and J-genes lead to similar distributions of loop lengths. To test this, we identified the closest germline genes for each antibody with ANARCI (27). We then group parent antibodies according to common V- and J-gene groups and compared the distributions of infilled loop lengths for each group (Figure 3E). While there may be some tendency for similar V- and J-genes to lead to similar distributions of infilled loop lengths, we observe considerable variation. This suggests that IgLM is not purely performing germline matching, but rather is considering other properties of the parent antibody.

#### Infilling generates diverse loop sequences

Diverse loop libraries are essential for discovering or optimizing sequences against an antigen target. To evaluate the diversity of infilled loops produced by IgLM, we measured the pairwise edit distance between each loop sequence and its closest neighbor amongst the sequences generated with the same sampling parameters. We then compared the diversity of sequences according to loop length and choice of sampling parameters (Figure 3F-G). Generally, we observe that generated loops are more diverse at longer lengths, as expected given the increased combinatorial complexity available as more residues are added. Increasing both sampling temperature and nucleus probability results in a greater diversity of sequences. However, these parameters affect the relationship between length and diversity in distinct ways. For a given loop length, increasing temperature produces more variance in the pairwise edit distance, while increases to nucleus probability provides a more consistent increase in diversity across loop lengths. Indeed, the marginal distribution of pairwise edit distance as nucleus probability is increased produces a much larger shift (Figure 3G, marginal) than that of temperature (Figure 3F, marginal). In practice, a combination of sampling parameters may be suitable for producing a balance of high-likelihood (low temperature and low nucleus probability) and diverse sequences sequences.

#### Infilled loops display improved developability

Developability encompasses a set physiochemical properties – including aggregation propensity and solubility – that are critical for the success of a therapeutic antibody. Libraries for antibody discovery or optimization that are enriched for sequences with improved developability can allieviate the need for time-consuming post-hoc engineering. To evaluate the developability of sequences produced by IgLM, we used high-throughput computational tools to calculate the aggregation propensity (SAP score (31)) and solubility (CamSol Intrinsic (32)) of the infilled therapeutic libraries. As a precursor to calculation of aggregation propensity, we used IgFold (21) to predict the structures of the infilled antibodies (including the unchanged light chains). We then compared the aggregation propensities and solubility values of the infilled sequences to those of the parent antibodies. For aggregation propensity, we observed a significant improvement (negative is better) by infilled sequences over the parent antibodies (Figure 4A, Figure S5). Similarly for solubility, infilled sequences tend to be more soluble than their parent antibodies (Figure 4B, Figure S6). In both cases, the largest improvements tend to correspond to the shorter loops. Further, we observe a positive correlation between improvements to aggregation propensity and solubility (Figure 4C, Figure S7). These results suggest that infilling can be used to generate libraries enriched for sequences with improved developability.

**Fig. 4.**
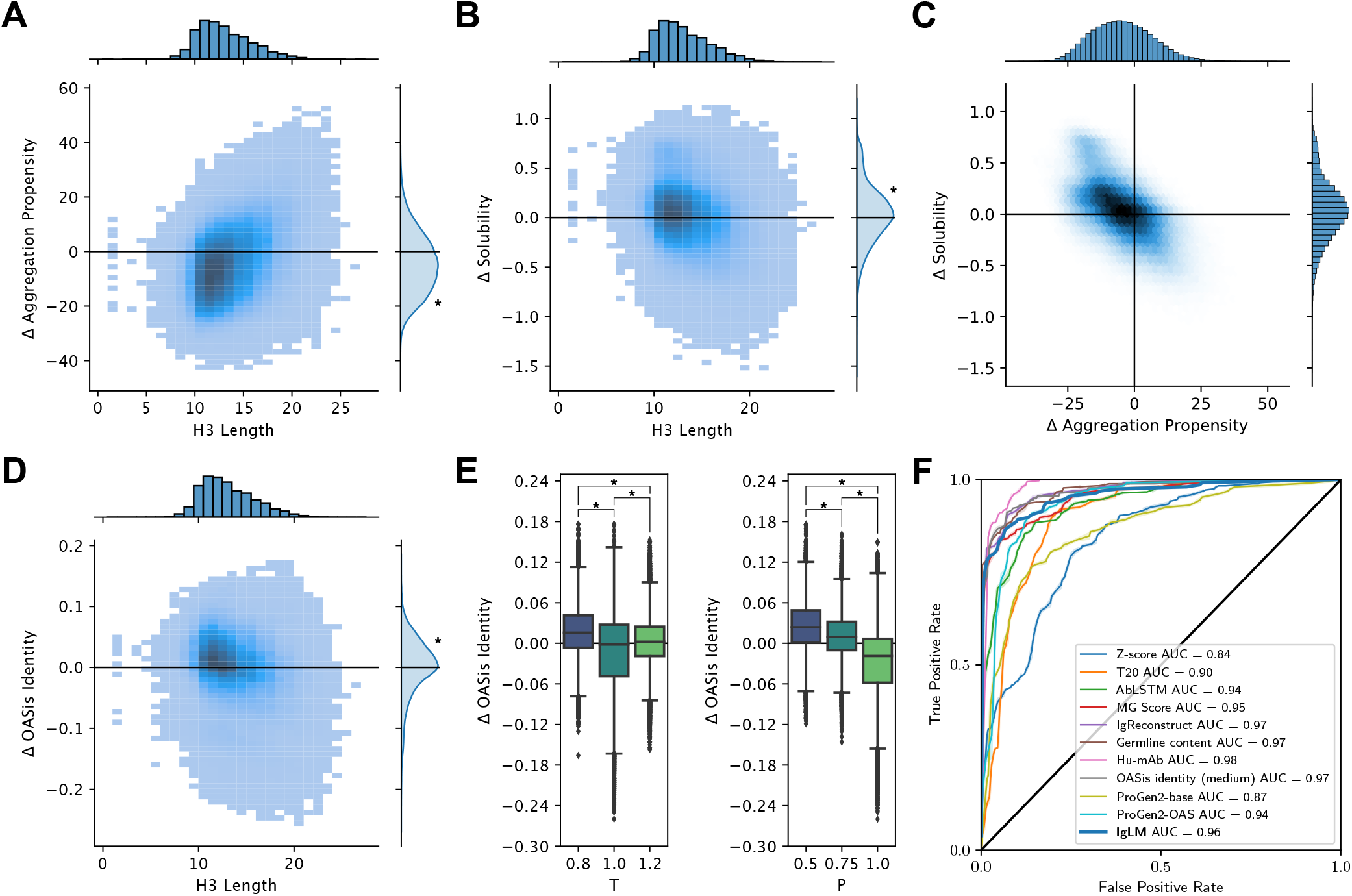
Therapeutic properties of infilled antibody libraries. Asterisks indicate statistical significance (p < 0.001) from a one-sample t-test (A, B, D) or a two-sample t-test (E). (A) Change in predicted aggregation propensity of infilled sequences relative to their parent antibodies. Infilled sequences display reduced aggregation propensity (negative is improved), particularly for shorter loops. (B) Change in predicted solubility of infilled sequences relative to their parent antibodies. Infilled sequences display increased solubility (positive is improved). (C) Relationship between predicted changes in aggregation propensity and solubility for infilled sequence libraries. (D) Change in humanness of infilled sequences relative to their parent antibodies. Humanness is calculated as the OASis identity of the heavy chain sequence, with positive larger values being more humanlike. (E) Relationship between sampling temperature (*T*) and nucleus probability (*P*) and change in human-likeness (OASis identity) of infilled heavy chains relative to their parent sequences. (F) Receiver operating characteristic (ROC) curves for human sequence classification methods. The area under the curve (AUC) is shown for each method.

We next investigated whether choice of sampling parameters affects the developability of infilled sequences. When we compared the aggregation propensity and solubility of infilled sequences according to the sampling temperature and nucleus sampling probability, we found marginal practical differences (Figure S8). This is likely explained by the relative consistency of infilled loop lengths across sampling parameters (Figure 3C). These results suggests that developability should not be a concern when tuning the diversity of a generated library.

#### Infilled loops are more human-like

Therapeutic antibodies must be human-like to avoid provoking an immune response and to be safe for use in humans. To evaluate the human-likeness of infilled sequences, we calculated the OASis identity (at medium stringency) (19). OASis divides an antibody sequence into a set of 9-mers and calculates the fraction that have been observed in human repertoires. Thus, higher OASis identity indicates a sequence that is more similar to those produced by humans. When compared to their respective parent antibodies, sequences infilled by IgLM were typically more human-like (Figure 4D). This is expected, given that IgLM is trained on natural human antibodies. We also investigated the impact of sampling parameters on the human-likeness of infilled sequences. For both sampling temperature and nucleus probability, we find that less restrictive sampling tends to produce less human-like sequences (Figure 4E). For practical purposes, this suggests that sampling with lower temperature and nucleus probability may be more suitable when immunogenicity is a concern.

### Sequence likelihood is an effective predictor of humanness

Likelihoods from autoregressive language models trained on proteins have been shown to be effective zero-shot predictors of protein fitness (15, 17). Antibody-specific language models in particular have been used to measure the “naturalness” of designed sequences (33), a measure related to humanness. To evaluate the effectiveness of IgLM for distinguishing human from non-human antibodies, we utilized the model’s likelihood to classify sequences from the IMGT mAb DB (34). Sequences in this set span a variety of species (human and mouse) and engineering strategies (e.g., humanized, chimeric, felinized). We considered all sequences not specifically labeled as human to be non-human, and calculated a likelihood (conditioned on human species) for each. All sequences had both a heavy and light chain, for which we calculated separate likelihoods and then multiplied.

We compared the performance of IgLM to that of a number of other methods previously benchmarked by Prihoda et al. (19) using a receiver operating characteristic (ROC) curve (Figure 4F). The results here for alternative methods are adapted from those presented by Prihoda et al, but with several redundant entries removed to avoid double-counting. We additionally evaluated model likelihoods from ProGen2-base and ProGen2-OAS (15), which are models similar to IgLM that contain significantly more parameters (764M). ProGen2-base is trained on a diverse set of protein sequences, while ProGen2-OAS is trained on a dataset similar to IgLM (OAS clustered at 85% sequence identity). We find that IgLM is competitive with state-of-the-art methods designed for human sequence classification, though not the best. Interestingly, IgLM outperforms ProGen2-OAS (ROC AUC of 0.96 for IgLM vs. 0.94 for ProGen2-OAS), despite having significantly fewer parameters (13M vs. 764M). This may result from the different strategies for constructing training datasets from OAS. By filtering at a less stringent 95% sequence identity, IgLM is likely exposed to a greater proportion of human antibody sequences, which dominate the OAS database. These distinctions highlight the importance of aligning training datasets with the intended application and suggest that training on only human sequences may further improve performance for human sequence classification.

## Discussion

Antibody libraries are a powerful tool for discovery and optimization of therapeutics. However, they are hindered by large fractions of non-viable sequences, poor developability, and immmunogenic risks. Generative language models offer a promising alternative to overcome these challenges through on-demand generation of high-quality sequences. However, previous work has focused entirely on contiguous sequence decoding (N-to-C or C-to-N) (15, 24). While useful, such models are not well-suited for generating antibody libraries, which vary in well-defined regions within the sequence, and for which changes may be undesirable in other positions. In this work, we presented IgLM, an antibody-specific language model for generation of full-length sequences and infilling of targeted residue spans. IgLM was trained for sequence infilling on 558M natural antibody sequences from six species. During training, we provide the model with conditioning tags that indicate the antibody’s chain type and species-of-origin, enabling controllable generation of desired types of sequences.

Concurrent work on autoregressive language models for antibody sequence generation have been trained on similar sets of natural antibody sequences and explored larger model sizes (15). However, models like ProGen2-OAS are limited in utility for antibody generation and design, as they are difficult to guide towards generation of specific types of sequences (e.g., species or chain types). Both this work and the ProGen2-OAS paper have utilized prompting strategies to guide model generation towards full-length sequences. While these strategies may help in some cases (particularly to overcome dataset limitations), significantly more reisdues may need to be provided to guide the model towards a specific sequence type (e.g., human vs rhesus heavy chain). In contrast, by including conditioning information for species and chain type in the model’s training, IgLM is able to generate sequences of the desired type without additional prompting. Still, as shown in this work, increasing the capacity of models like IgLM may lead to better performance for sequence infilling (lower perplexity) and scoring (better likelihood estimation), a promising direction for future work.

IgLM’s primary innovation is the ability to generate infilled residue spans at specified positions within the antibody sequence. In contrast to traditional generative language models that only consider preceding the residues, this enables IgLM to generate within the full context of region to be infilled. We demonstate the utility of infilling by generating libraries for 49 therapeutic antibodies. We found that IgLM was capable of generating diverse CDR H3 loop sequences, and that diversity was largely tunable by choice of sampling parameters. Further, the infilled libraries possessed desirable developability traits (aggregation propensity, solubility) while being more human-like on average than their parent sequences. Notably, IgLM achieves these improvements over antibodies that are already highly optimized, as all of the parent sequences have been engineered for mass-production and use in humans. Although we focused on antibody loop infilling in this work, similar strategies may be useful for proteins generally. For example, a universal protein sequence infilling model may be applicable to redesign of contiguous protein active sites or for generating linkers between separate domains for protein engineering.

## Code and Data Availability

Code and pre-trained models for IgLM are available at https://github.com/Graylab/IgLM. All generated sequences and predicted structures will be deposited to Zenodo upon publication.

## Methods

### Infilling formulation

Designing spans of amino acids within an antibody sequence can be formulated as an infilling task, similar to text-infilling in natural langauge. We denote an antibody sequence *A* = (*a*_1_, … *a_n_*), where *a_i_* represents the amino acid at position *i* of the antibody sequence. To design a span of length *m* starting at position *j* along the sequence, we first replace the span of amino acids *S* = (*a_j_*, … *a*_*j*+*m*–1_) with a single [`MASK`] token to form a sequence *A*_\*S*_ = (*a*_1_, … *a*_*j*–1_; [`MASK`], *a*_*j*+*m*_, … *a_n_*). To generate reasonable variable-length spans to replace *S* given *A*_\*S*_, we seek to learn a distribution *p*(*S*|*A*_\*S*_).

We draw inspiration from the Infilling by Language Modeling (ILM) framework proposed for natural language infilling (25) to learn *p*(*S*|*A*_\*S*_). For assembling the model input, we first choose a span *S* and concatenate *A*_\*S*_, [`SEP`], *S*, and [`ANS`]. We additionally prepend conditioning tags *c_c_* and *c_s_* to specific the chain type (heavy or light) and species-of-origin (e.g., human, mouse, etc.) of the antibody sequence. The fully formed sequence of tokens **X** for IgLM is:

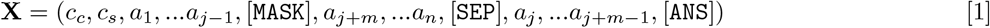

We then train a generative model with parameters *θ* to maximize *p*(**X**|*θ*), which can be decomposed into a product of conditional probabilities:

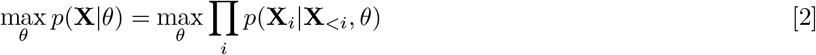

### Model implementation

The IgLM model uses a modified version of the GPT-2 Transformer decoder architecture (35) as implemented in the HuggingFace Transformers library (36). We trained two models, IgLM and IgLM-S, for sequence infilling. Hyperparameter details are provided in Table 1.

**Table 1.** IgLM model hyperparameters.

|  | IgLM | IgLM-S |
| --- | --- | --- |
| Number of layers | 4 | 3 |
| Embedding dimension | 512 | 192 |
| Hidden dimension | 512 | 192 |
| Attention heads | 8 | 6 |
| Feed-forward dimension | 2048 | 768 |
| Total parameters | 12,889,600 | 1,439,616 |

### Antibody sequence dataset

To train IgLM, we collected unpaired antibody sequences rom the Observed Antibody Space (OAS) (18). OAS is a curated set of over one billion unique antibody sequences compiled from over eighty immune repertoire sequencing studies. After removing sequences indicated to have potential sequencing errors, we were left with 809M unique antibody sequences. We then clustered these sequences using LinClust (37) at 95% sequence identity, leaving 588M non-redundant sequences. The distribution of sequences corresponding to each species and chain type are documented in Figure 1B and Table S1. The dataset is heavily skewed towards human antibodies, particularly heavy chains, which make up 70% of all sequences. We held out 5% of sequences as a test set to evaluate model performance. Of the remaining sequences, we used 558M sequences for training and 1M for validation.

### Model training

During training, for each sequence *A* = (*a*_1_,…, *a_n_*) we chose a mask length *m* uniformly at random from [10, 20] and a starting position *j* uniformly at random from [1, *n* – *m* +1]. We prepended two conditioning tags *c_c_* and *c_s_* denoting the chain type and species-of-origin of each sequence as annotated in the OAS database. Models were trained with a batch size of 512 and 2 gradient accumulation steps using DeepSpeed (38, 39). Training required approximately 3 days when distributed across 4 NVIDIA A100 GPUs.

## ACKNOWLEDGMENTS

We thank Dr. Sai Pooja Mahajan and Dr. Rahel Frick for insightful discussions and advice. This work was supported by the National Science Foundation grant DBI-1950697 (R.W.S.) and National Institutes of Health grants R01-GM078221 (J.A.R.) and R35-GM141881 (J.A.R.) J.A.R. was supported as a Johns Hopkins-AstraZeneca Fellow. Computation was performed using the Advanced Research Computing at Hopkins (ARCH) core facility.

## Notes

### Competing Interest Statement

The Johns Hopkins University has filed one or more patent application(s) related to this technology. R.W.S., J.A.R., and J.J.G. are named as inventors on these application(s).

### Summary of Updates

Expanded analysis of controllable generation and therapeutic antibody library generation.

