## Supplemental materials for "Generative language modeling for antibody design"

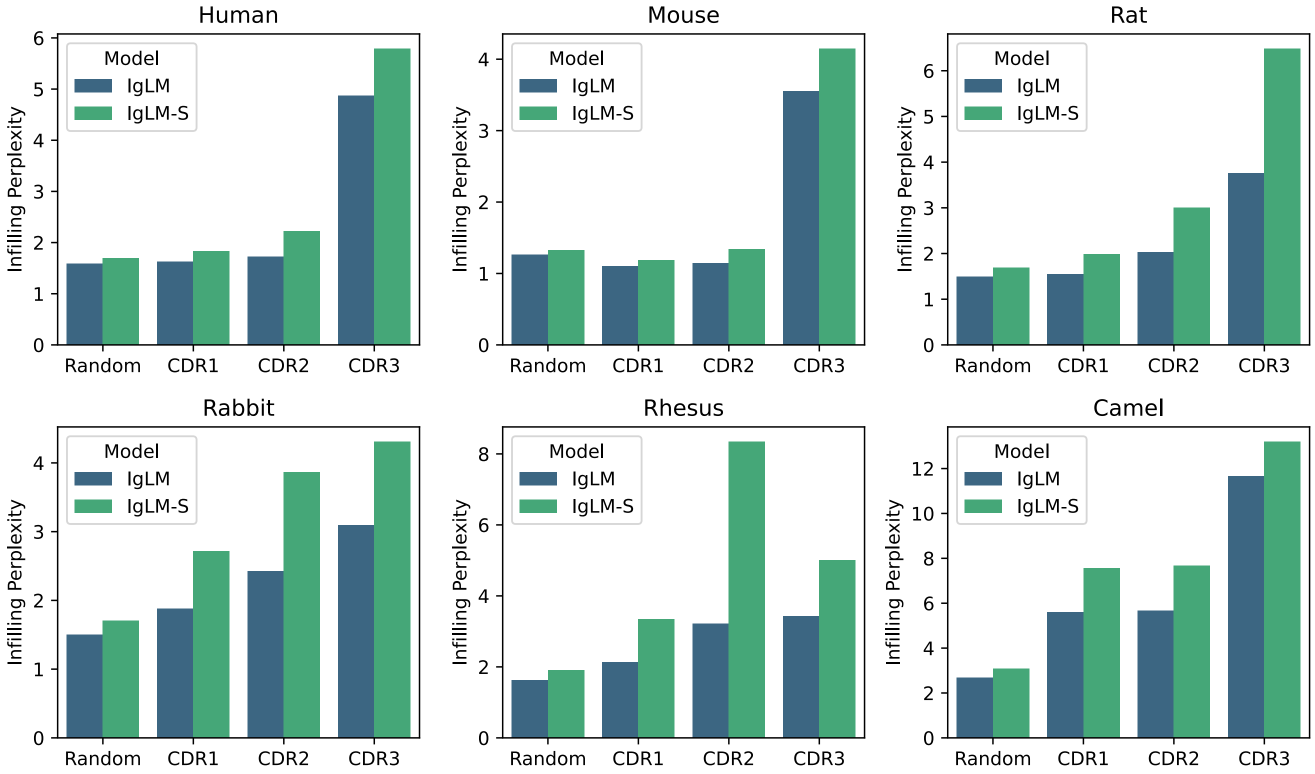


**Fig. S1.** Infilling perplexity for IgLM and IgLM-S on heldout test dataset of 30M sequences, divided by species-of-origin. Values are reported for CDR loops and random spans of 10-20 residues within sequences.

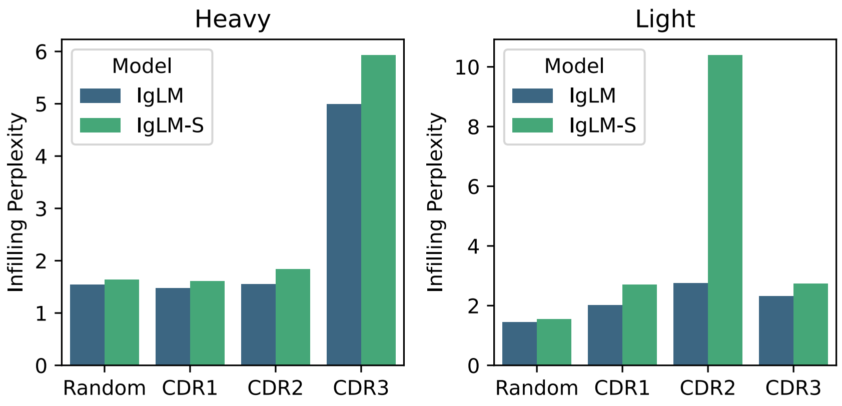


**Fig. S2.** Infilling perplexity for IgLM and IgLM-S on heldout test dataset of 30M sequences, divided by chain type. Values are reported for CDR loops and random spans of 10-20 residues within sequences.

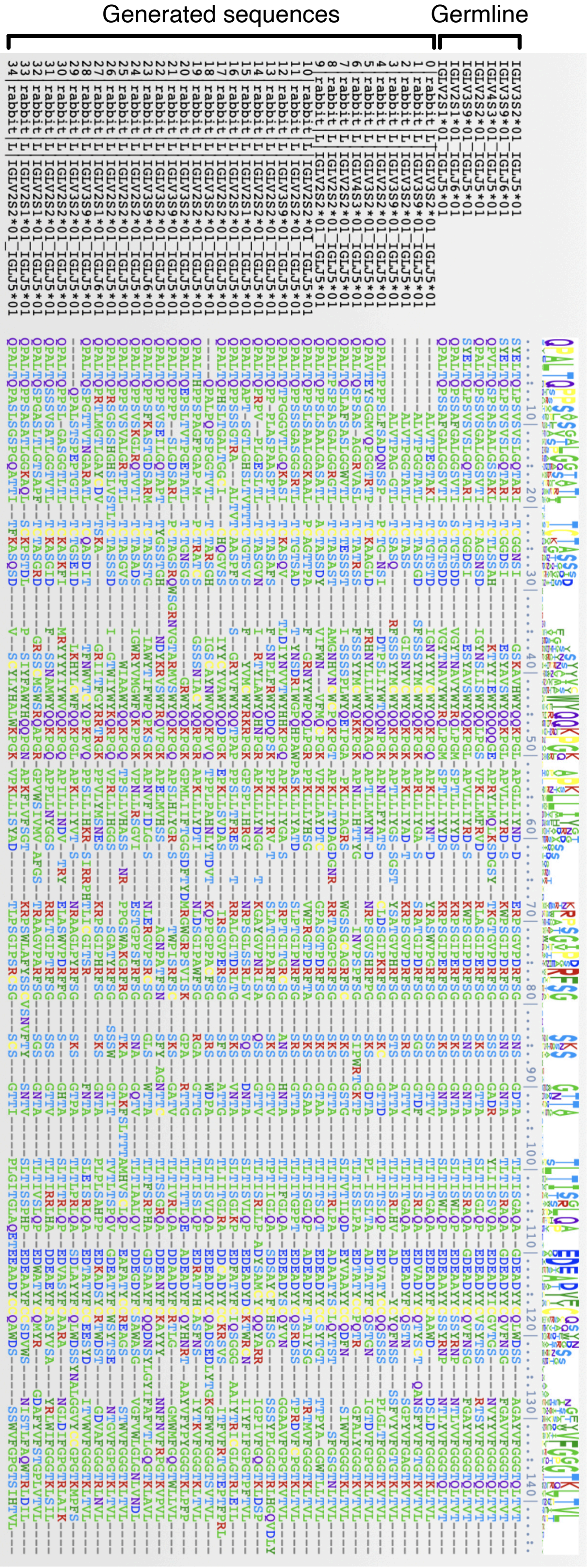


**Fig. S3.** Alignment of generated rabbit light chain sequences with the closest germline sequences assigned by ANARCI. Sequences were aligned with Clustal-Omega.

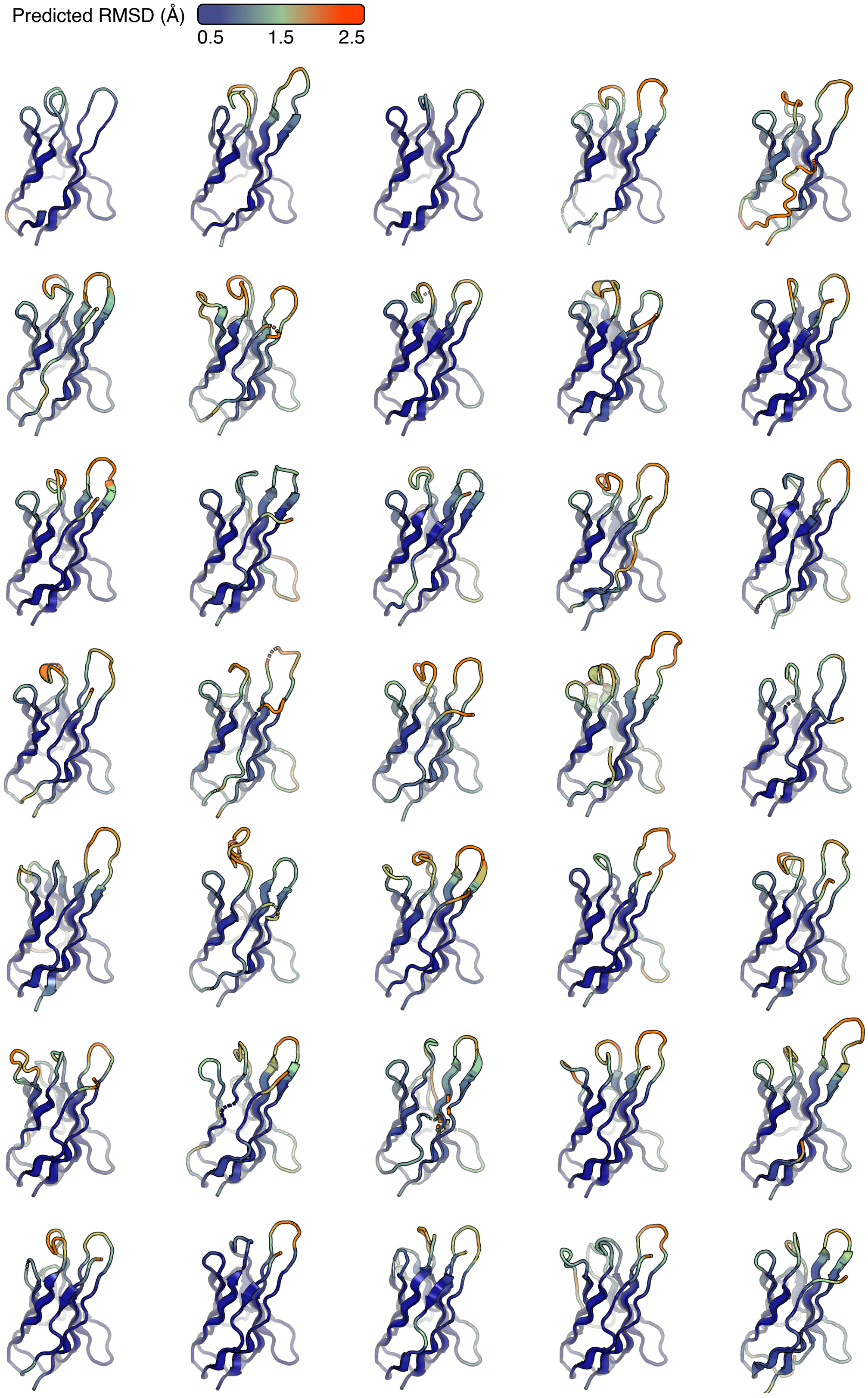


**Fig. S4.** Prediction of generated rabbit light chain sequences by IgFold. Structures are colored by predicted RMSD.

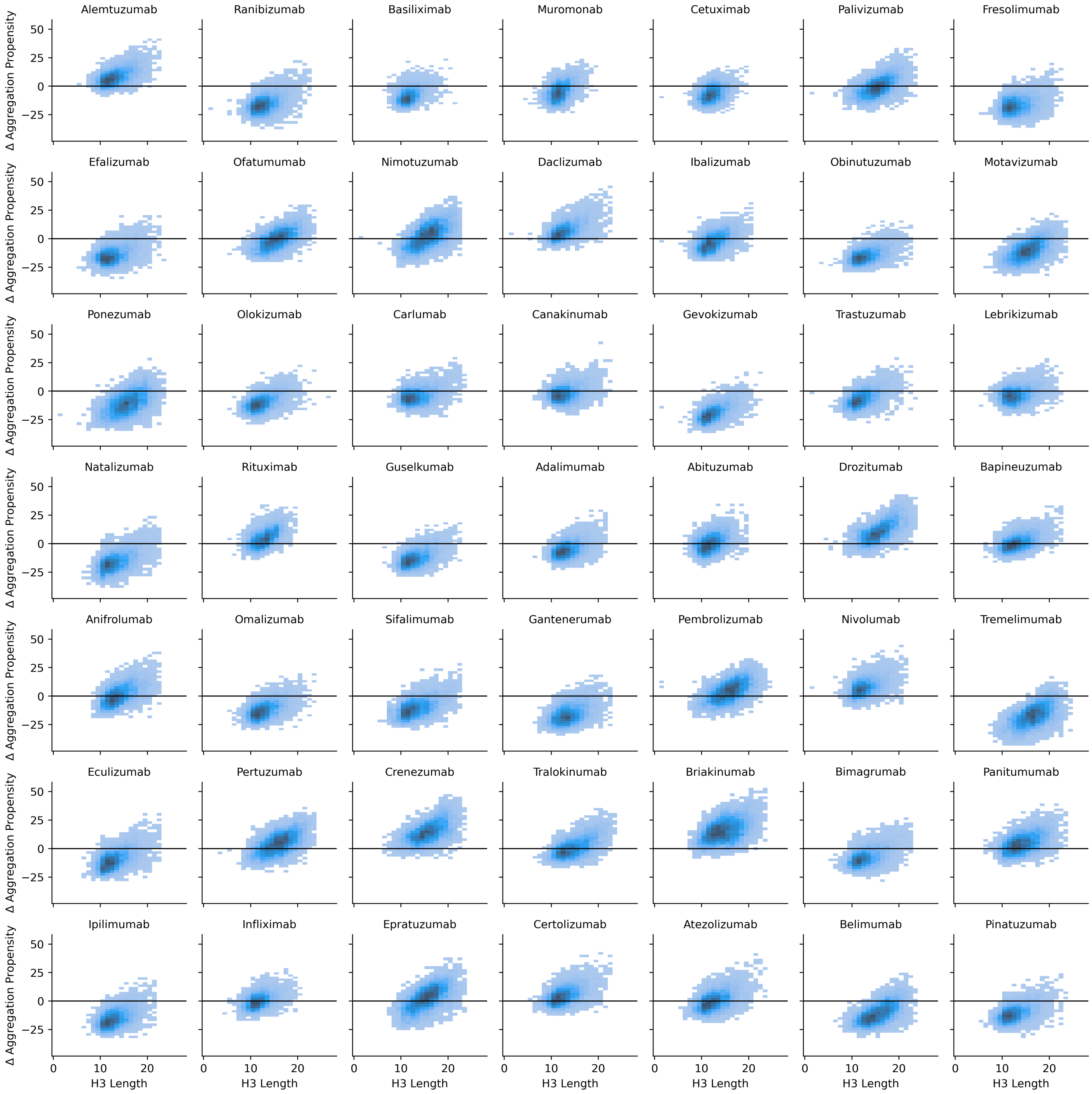


**Fig. S5.** Comparison of infilled CDR H3 loop length and change in aggregation propensity (relative to parent, lower is better) for individual infilled libraries.

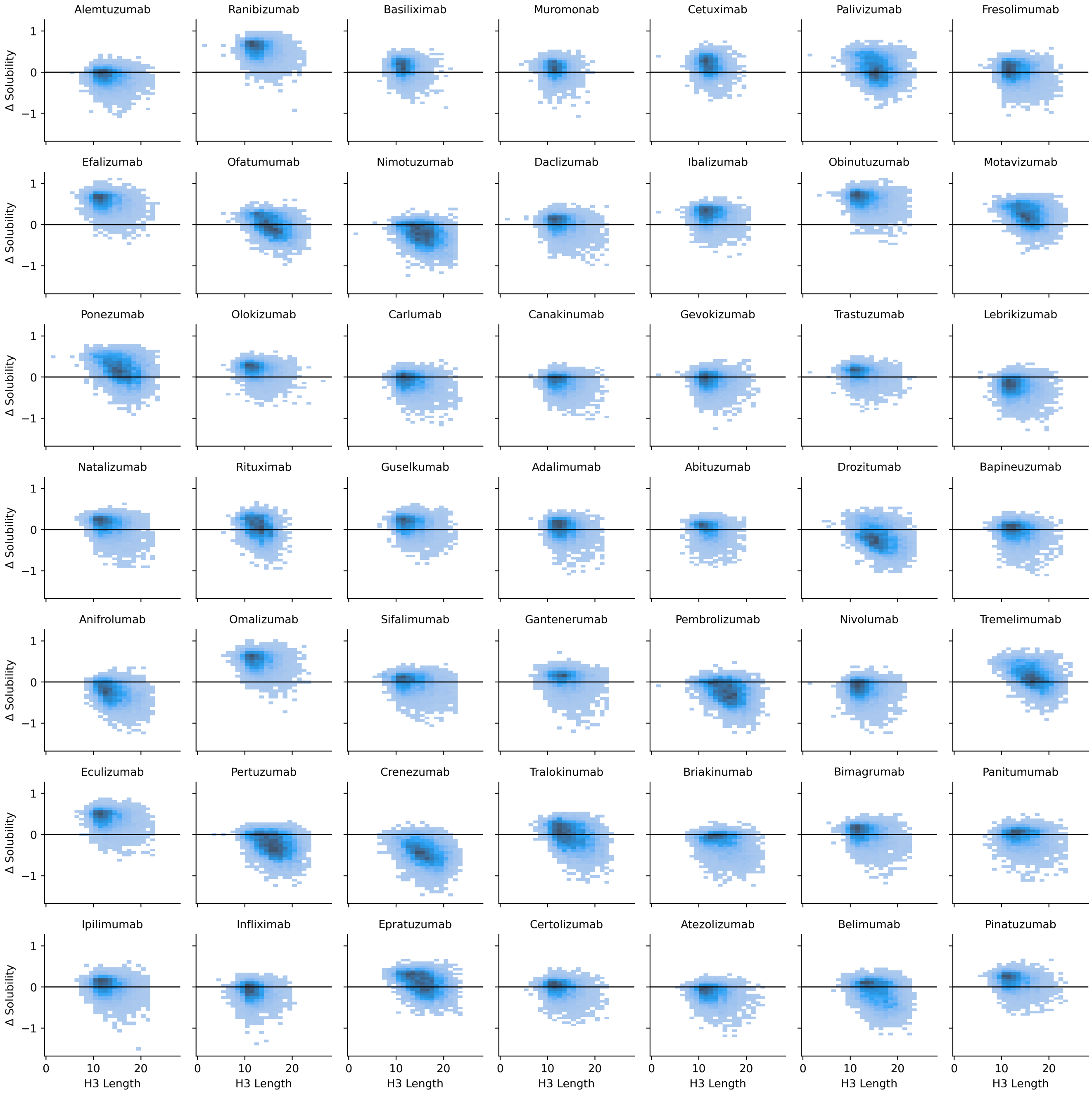


**Fig. S6.** Comparison of infilled CDR H3 loop length and change in solubility (relative to parent, higher is better) for individual infilled libraries.

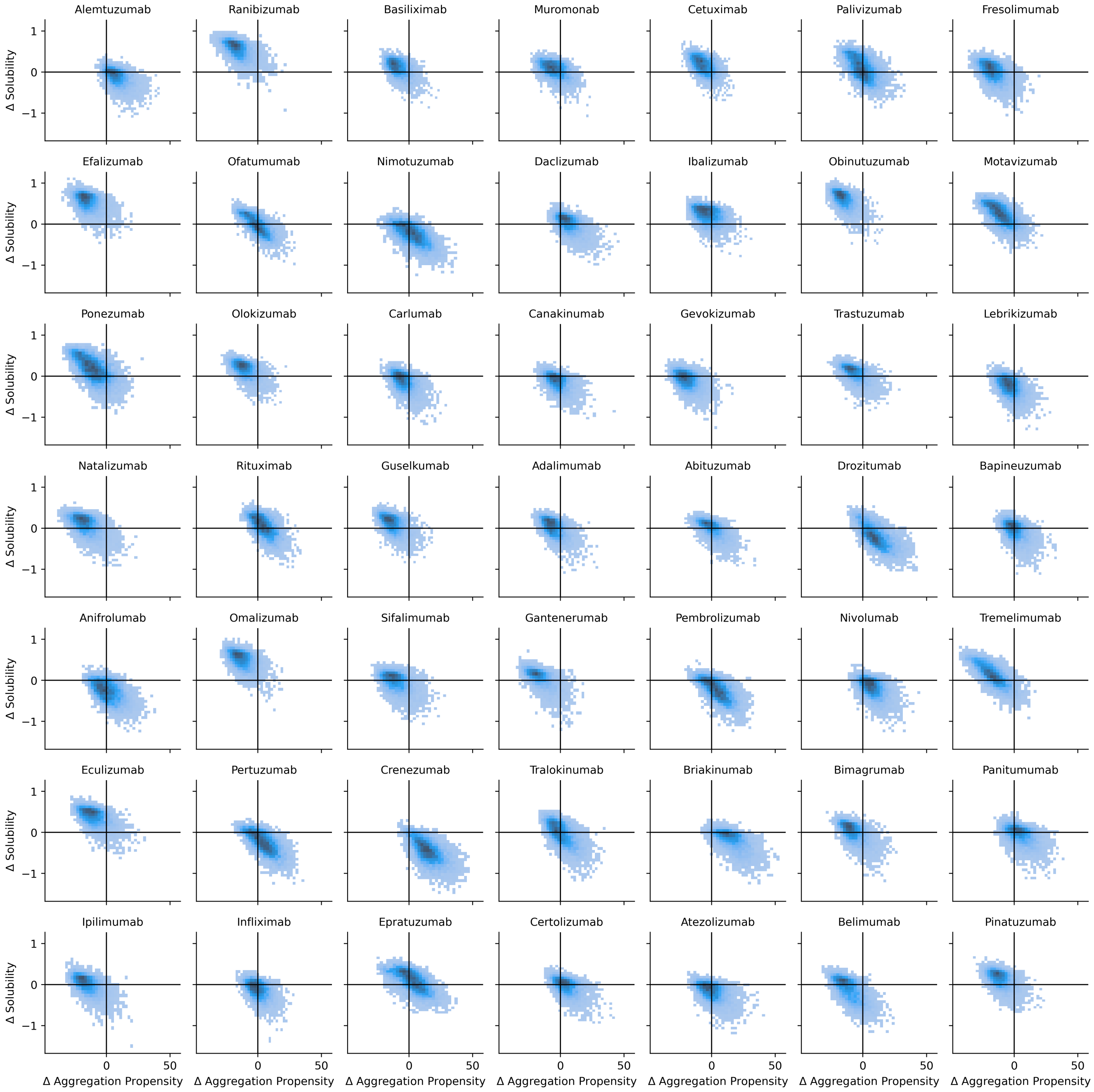


**Fig. S7.** Relationship between changes in aggregation propensity and solubility for individual infilled libraries.

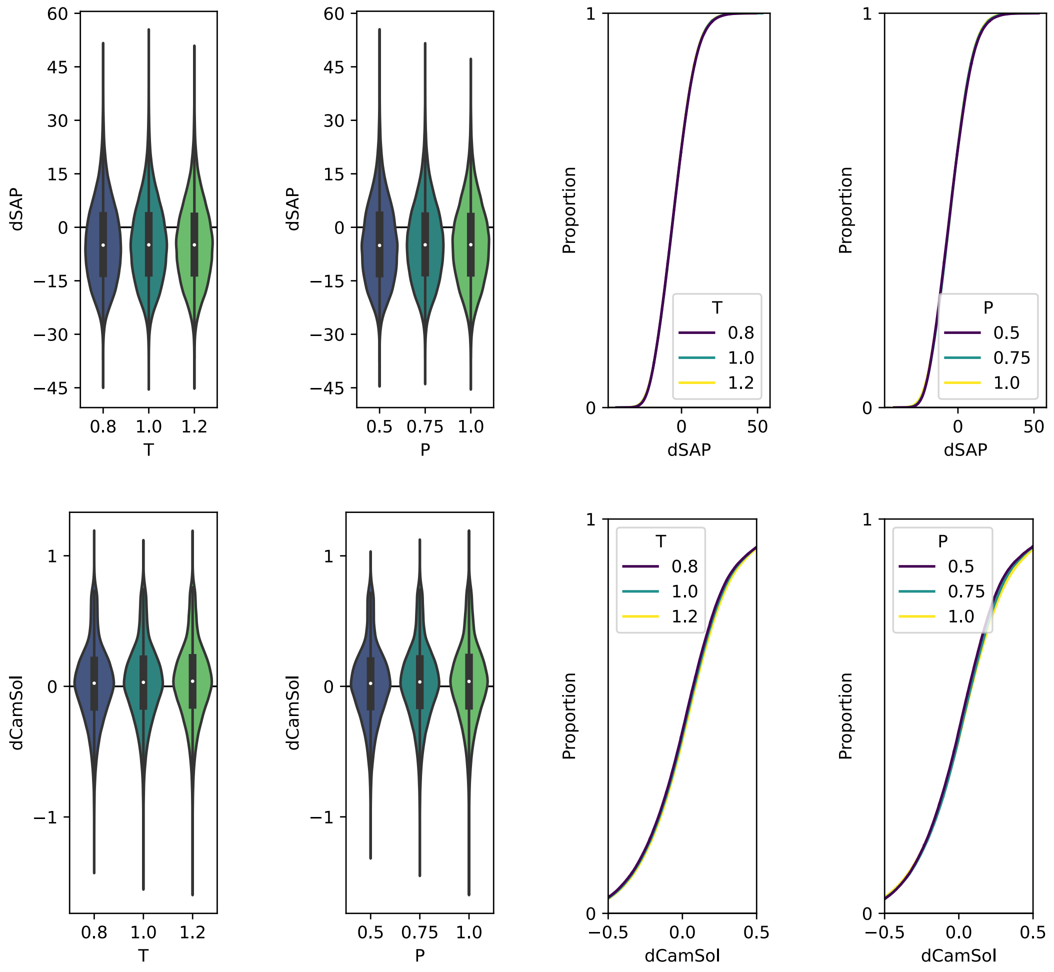


**Fig. S8.** Impact of sampling parameters on developability of infilled libraries. Library properties are largely unaffected by choice of sampling temperature ($T$) and nucleus sampling probability ($P$).

**Table S1.** Distribution of sequences in clustered OAS dataset.

| Species | Heavy Chains | Light Chains | Total |
|---|---|---|---|
| Human | 412,807,447 | 70,584,881 | 483,392,328 |
| Mouse | 93,360,086 | 3,198,407 | 96,558,493 |
| Camel | 1,091,641 | - | 1,091,641 |
| Rat | 3,700,086 | 0 | 3,700,086 |
| Rabbit | 2,644,903 | 0 | 2,644,903 |
| Rhesus | 381,021 | 719,674 | 1,100,695 |
| **Total** | **513,985,184** | **74,502,962** | **588,488,146** |

**Table S2.** Full-length sequence generation parameters.

| Description | Chain token | Species token | Initial residues | Number generated |
|---|---|---|---|---|
| Human heavy chain | [HEAVY] | [HUMAN] | EVQ | 20,000 |
| Human light chain (lambda) | [LIGHT] | [HUMAN] | QSA | 10,000 |
| Human light chain (kappa) | [LIGHT] | [HUMAN] | DIQ | 10,000 |
| Mouse heavy chain | [HEAVY] | [MOUSE] | QVQ | 20,000 |
| Mouse light chain (lambda) | [LIGHT] | [MOUSE] | QAV | 10,000 |
| Mouse light chain (kappa) | [LIGHT] | [MOUSE] | DIV | 10,000 |
| Camel heavy chain | [HEAVY] | [CAMEL] | QVQ | 20,000 |
| Rabbit heavy chain | [HEAVY] | [RABBIT] | QEQ | 20,000 |
| Rabbit light chain (lambda) | [LIGHT] | [RABBIT] | QPA | 10,000 |
| Rabbit light chain (kappa) | [LIGHT] | [RABBIT] | ALV | 10,000 |
| Rat heavy chain | [HEAVY] | [RAT] | EVQ | 20,000 |
| Rat light chain (lambda) | [LIGHT] | [RAT] | QAV | 10,000 |
| Rat light chain (kappa) | [LIGHT] | [RAT] | GIQ | 10,000 |
| Rhesus heavy chain | [HEAVY] | [RHESUS] | QVQ | 20,000 |
| Rhesus light chain (lambda) | [LIGHT] | [RHESUS] | QSV | 10,000 |
| Rhesus light chain (kappa) | [LIGHT] | [RHESUS] | AIQ | 10,000 |
| **Total** | | | | **220,000** |

**Table S3.** Adherence to species conditioning tags for full-length generation. Percentage of matches between conditioning tag used for generation and species classification from ANARCI for each generation configuration.

| Chain token | Species token | T = 0.6 | T = 0.8 | T = 1.0 | T = 1.2 | Overall |
|---|---|---|---|---|---|---|
| [HEAVY] | [HUMAN] | 99.98% | 99.94% | 99.66% | 99.30% | 99.72% |
| [HEAVY] | [MOUSE] | 99.94% | 99.34% | 98.32% | 96.62% | 98.55% |
| [HEAVY] | [CAMEL] | 98.36% | 97.72% | 96.76% | 92.72% | 96.39% |
| [HEAVY] | [RABBIT] | 100.00% | 100.00% | 99.96% | 99.98% | 99.98% |
| [HEAVY] | [RAT] | 0.00% | 0.00% | 0.00% | 0.00% | 0.00% |
| [HEAVY] | [RHESUS] | 99.00% | 94.82% | 87.14% | 71.58% | 88.14% |
| [LIGHT] | [HUMAN] | 100.00% | 99.98% | 99.92% | 99.40% | 99.82% |
| [LIGHT] | [MOUSE] | 13.84% | 86.62% | 16.64% | 22.46% | 34.89% |
| [LIGHT] | [RABBIT] | 14.56% | 5.84% | 1.37% | 2.22% | 6.89% |
| [LIGHT] | [RAT] | 0.00% | 0.00% | 0.02% | 0.15% | 0.04% |
| [LIGHT] | [RHESUS] | 70.14% | 49.02% | 44.80% | 51.15% | 54.03% |
